# Genomic consequences of a recent three-way admixture in supplemented wild brown trout populations revealed by ancestry tracts

**DOI:** 10.1101/302380

**Authors:** Maeva Leitwein, Pierre-Alexandre Gagnaire, Erick Desmarais, Patrick Berrebi, Bruno Guinand

## Abstract

Understanding the evolutionary consequences of human-mediated introductions of domestic strains into the wild and their subsequent admixture with natural populations is of major concern in conservation biology. In the brown trout *Salmo trutta*, decades of stocking practices have profoundly impacted the genetic makeup of wild populations. Small local Mediterranean populations in the Orb River watershed (Southern France) have been subject to successive introductions of domestic strains derived from the Atlantic and Mediterranean lineages. However, the genomic impacts of two distinct sources of stocking (locally-derived vs divergent) on the genetic integrity of wild populations remain poorly understood. Here, we evaluate the extent of admixture from both domestic strains within three wild populations of this watershed, using 75,684 mapped SNPs obtained from double-digest restriction-site-associated DNA sequencing (dd-RADseq). Using a local ancestry inference approach, we provide a detailed picture of admixture patterns across the brown trout genome at the haplotype level. By analysing the chromosomal ancestry profiles of admixed individuals, we reveal a wider diversity of hybrid and introgressed genotypes than estimated using classical methods for inferring ancestry and hybrid pedigree. In addition, the length distribution of introgressed tracts retained different timings of introgression between the two domestic strains. We finally reveal opposite consequences of admixture on the level of polymorphism of the recipient populations between domestic strains. Our study illustrates the potential of using the information contained in the genomic mosaic of ancestry tracts in combination with classical methods based on allele frequencies for analysing multiple-way admixture with population genomic data.

## Introduction

From the early 1950’s, hybridization has been associated to intentional or unintentional disturbance of natural systems by human activities (Anderson 1948, 1949, 1953; Anderson and Stebbins 1954; Wiegand 1935). Since then, human-mediated hybridization (HMH) between source and recipient populations has become an increasingly significant eco-evolutionary concern in conservation biology (Levin *et al.*, 1996; Rhymer and Simberloff, 1996). Despite the continued technical improvements in the analysis of hybridization and admixture, the debate about the costs and benefits of HMH is still ongoing (e.g., Bohling, 2016; Wayne and Shaffer, 2016; Grabenstein and Taylor, 2018). On the one hand, hybridization can be a source of beneficial evolutionary novelties through adaptive introgression (Anderson, 1949; Stebbins, 1950; Anderson and Stebbins, 1954; Heiser, 1951; Whitney *et al.*, 2015; Schumer *et al.*, 2014; Yakimowski and Rieseberg, 2014; Lamichhaney *et al.*, 2017; Runemark *et al.*, 2018). On the other hand, hybridization can erase diversity resulting from speciation and natural selection, leading to genetic swamping and loss of local adaptation (e.g. Laikre *et al.*, 2010; Randi, 2008; Rhymer and Simberloff, 1996; Todesco *et al.*, 2016). The emerging view offered by genome-wide surveys is that HMH often results in a mix of positive and negative effects that are scattered across the genome, leading to the “haplotype block view” of hybridization introducing adaptive and maladaptive “*blocks of genic materials belonging to different adaptive systems*” (Anderson and Stebbins 1954).

This ‘genic block’ metaphor referred to what is now coined as local ‘ancestry tracts’ resulting from multigenerational gene flow and recombination events among hybridizing taxa (Buerkle and Lexer, 2008; Gravel, 2012; Gompert and Buerkle, 2013; Liu *et al.*, 2013; Liang and Nielsen, 2014). The identification of ancestry tracts is based on the analysis of marker associations in admixed individuals with the goal of recovering continuous ancestry blocks inherited from ancestral taxa, sometimes including explicitly linkage information (e.g. Price *et al.*, 2009; Lawson *et al.*, 2012; Loh *et al.*, 2013; Zhou *et al.*, 2017). The number and the length distribution of such tracts are informative (i) on the admixture level and variation at both a global (i.e. the relative ancestry proportions of each admixed individual; Buerkle, 2005) and local genomic scale (or locus-specific; i.e. the ancestry of each haplotype at a particular locus; Falush *et al.*, 2003), but also (ii) on the number of generations since initial admixture because recombination progressively breaks down ancestry tracts across generations (e.g. Pool and Nielsen, 2009; Gravel, 2012; Corbett-Detig and Nielsen, 2017).

The use of ancestry tracts has helped understanding the history of admixture during the recent and past history of human populations, including admixture between modern human and extinct archaic lineages (e.g., Wall *et al.*, 2013; Racimo *et al.*, 2015). More recently, these approaches have been also applied in domesticated plant and animal species to characterize the nature, origin and timing of introgression (e.g. VonHoldt *et al.*, 2016; Hufford *et al.*, 2013; Flori *et al.*, 2014; Galaverni *et al.*, 2017; Nelson *et al.*, 2017; Medugorac *et al.*, 2017; Wang *et al.*, 2017). The development of next-generation sequencing (NGS) technologies together with statistical development and associated softwares (a review in Johnson *et al.*, 2015) has offered an unprecedented way to study admixture and introgression in many hybridizing taxa including both model and non-model species (Kohn *et al.*, 2006; Primmer, 2009; Allendorf *et al.*, 2010; Ouborg *et al.*, 2010; Angeloni *et al.*, 2012; Narum *et al.*, 2013; Steiner *et al.*, 2013; Hoffmann *et al.*, 2015). However, the use of local ancestry blocks to characterize the genomic landscape of admixture remains scarce in conservation genomic studies (but see VonHoldt *et al.*, 2016; Galaverni *et al.*, 2017; Duranton *et al.*, 2017).

Because of their important socio-economic status, salmonids are amongst the most worldwide anthropized fish families (Antunes *et al.*, 1999; Fraser, 2008; Davidson *et al*., 2010). Stocking practices and enhancement using hatchery fishes are common in Salmonids (Antunes *et al.*, 1999; Aprahamian *et al.*, 2003; Sundt-Hansen *et al.*, 2015) and often result in gene flow between domestic and wild populations, which becomes a central issue to many conservation programs (Waples, 1991; Naish *et al.*, 2007; Fraser, 2008; Harbicht *et al.*, 2014; Skaala *et al.*, 2014; Glover *et al.*, 2017). The impacts of HMH on neutral genetic diversity (e.g. Laikre *et al.* 2010; Fernández-Cebrián *et al.* 2014), adaptation (e.g. McGinnity *et al*., 2003; Muhlfeld *et al.*, 2009; Le Cam *et al.*, 2015) and genetic integrity of recipient populations (e.g. Eldridge *et al.*, 2009; Valiquette *et al.* 2014; Ozerov *et al.*, 2016) have been assessed using molecular markers, but without using linkage information. Now that extensive genomic resources are available including dense linkage maps (e.g. Lien *et al.*, 2011; Gagnaire *et al.*, 2013; Gonen *et al.*, 2014; Brieuc *et al.*, 2014; McKinney *et al.*, 2015; Sutherland *et al.*, 2016; Tsai *et al.*, 2016; Leitwein *et al.*, 2017) and reference genomes (Berthelot *et al.*, 2014; Lien *et al.*, 2016), it becomes possible to explore the potential of local ancestry blocks for studying the genomic consequences of HMH in salmonids.

The brown trout (*Salmo trutta* L.) is a widely distributed Eurasian species with a complex evolutionary history (Elliott, 1994; Bernatchez and Osinov, 1995; Sanz, 2017). The species has been heavily impacted by human activities and is subject to stocking practices that are known to genetically impact wild populations to various degrees (e.g. Poteaux *et al*., 1999; Berrebi *et al.*, 2000; Almodóvar *et al.*, 2001; Hansen, 2002; Hansen *et al.*, 2009). In France, the Atlantic hatchery lineage which has been domesticated for decades (Bohling *et al.*, 2016) has been largely used for restocking and enhancement of wild Mediterranean local populations that belong to a distinct evolutionary lineage (e.g. Bernatchez 2001). In order to avoid mixing different lineages by supplementing with the domestic Atlantic strain, several recent attempts have been made to develop local domestic Mediterranean strains with the aim to release locally adapted genotypes into the wild (Bohling *et al.* 2016). However, the impacts of these different stocking practices have not been assessed at the genomic level yet.

The aim of this study was to investigate admixture from a distantly (i.e. Atlantic) and a closely (i.e. Mediterranean) related hatchery source strain into wild Mediterranean brown trout populations. We assessed the mosaic of introgressed tracts for characterizing the complex makeup of admixed genotypes within three wild brown trout populations, and improve the distinction between ‘early-generation hybrids’ such as F_1_, F_2_ and backcrosses, and ‘late-generation hybrids’ produced by several generations of admixture in the wild. Finally, the relative timing of admixture was estimated from both domestic source populations within each of the three wild populations. We took advantage of the high-density linkage map recently developed by Leitwein *et al.* (2017) to address these issues using RAD markers.

## Materials and methods

### Sampling

Wild Mediterranean brown trout were sampled from three tributaries of the Orb River watershed in southern France (April-May 2015). Eighty-two individuals were caught by electrofishing, fin clipped and released under the supervision of the Hérault French Fishing Federation. The study includes a total of 23 individuals from the upper Orb River, 45 from the Gravezon River, and 14 from the Mare River (Table 1). Farmed fish (N = 102) were sampled in 2014 at the Babeau hatchery to genetically characterize each of the two domestic strains that were used for stocking. They consisted of 61 individuals from the Atlantic domestic strain, which is commonly used for stocking practices in France for decades (Bohling *et al.* 2016), and 41 individuals from a Mediterranean domestic strain, which was initially developed by the Hérault French Fishing Federation in 2004 (Table 1). The Mediterranean domestic strain was founded exclusively with individuals from the Gravezon River (Hérault French Fishing Federation, *pers. comm.*). Additional information on the stocking of Atlantic brown trout in France and sampling are provided in Bohling *et al.* (2016) and Leitwein *et al.* (2016), respectively. Sixty individuals (including both farmed and wild) of the 184 analyzed in this study were part of a previous study describing genome-wide patterns of nucleotide diversity in Atlantic and Mediterranean brown trout lineages using dd-RAD-seq (Leitwein *et al.* 2016).

**Table 1.** Number of individuals sampled for the hatchery strain and wild population *N*) along with the number of individuals after filtering for missing data (*Nfiltered*) and the number of individuals considered as pure domestic or pure Mid with ADMIXTURE (*Npure*) see text for details.

| Samples | labels | *N* | *N<sub>filtered</sub>* | *N<sub>pure</sub>* |
| --- | --- | --- | --- | --- |
| Hatchery strain |  |  |  |  |
| Cauteret<br>(Atlantic) | ATL | 61 | 54 | 54 |
| Babeau<br>(Mediterranean) | domMED | 41 | 40 | 40 |
| River Orb |  |  |  |  |
| La Mare | Mare | 14 | 14 | 4 |
| Gravezon | Grav | 45 | 45 | 15 |
| Upper Orb | Orb | 23 | 23 | 10 |
| Total |  | 184 | 176 | 123 |

### DNA extraction and sequencing

Individual genomic DNA was extracted from caudal fin clips using the commercial KingFisher Flex Cell and Tissues DNA Kit. DNA quantity and quality were evaluated using both NanoDrop ND-8000 spectrophotometer (Thermo Fisher Scientific) and Qubit 1.0 Fluorometer (Introgen, Thermo Fisher Scientific). The double digested restriction site-associated DNA (dd-RAD) library preparation protocol followed Leitwein *et al.* (2016). Briefly, the two restriction enzymes *Eco*RI-HF and *MspI* were used to digest individual genomic DNA, which was subsequently submitted to adaptor ligation (one with unique barcodes for each individual and one common to 48 individuals containing Illumina index). Each library consisting of 48 individuals with unique barcodes pooled in equimolar proportions was fragmented and submitted to size selection to retain fragments ranging from 200 to 700bp using CleanPCR beads. The library was then amplified by PCR and sequenced with Illumina HiSeq2500, producing 125bp paired-end reads.

### Genotyping

The bioinformatics pipeline used for SNP calling has been described in Leitwein *et al.* (2016). Briefly, reads where demultiplexed, cleaned and trimmed to 120bp using process_radtags.pl implemented in STACKS v1.35 (Catchen *et al.*, 2013). Individual genotypes at RAD markers were determined using a reference mapping approach with the Atlantic salmon genome taken as a reference (GenBank accession number: GCA_000233375.4_ICSASG_v2; Lien *et al.*, 2016). The `BWA_mem` program v. 0.7.9 (Li and Durbin, 2010) was used to align reads along *S. salar* genome before calling SNPs for each individual with the `pstacks` module (using *m* = 3 and the bounded error model with α = 0.05). We used a previously established reference RAD catalogue constructed by Leitwein *et al.* (2016, 2017) (--`catalog` in the `cstacks` module). Each of the 184 individuals was then matched against this catalogue with `sstacks`. Seven domestic Atlantic and one domestic Mediterranean individuals were removed because of high percentage of missing data (>20%), resulting in a final data set of 176 individuals (82 wild Mediterranean individuals, 54 Atlantic and 40 Mediterranean domestic individuals). Finally, the `populations` module was used to generate a genotype dataset in VCF and PLINK formats containing loci passing the following filters: (*i*) a minimum stacks depth of 5 reads; (*ii*) a genotype call rate of at least 80% within each of the five populations, (*iii*) a minimum allele frequency of 2%, and (*iv*) a maximum observed heterozygosity of 60%. A single representative of each overlapping site was kept with the option --`ordered_export` in STACKS.

### Inference of admixture and hybridization

We used the filtered dataset containing 86,175 SNPs to estimate individual cluster membership using ADMIXTURE v1.3 (Alexander *et al.*, 2009). This program provides an estimation of individual ancestry proportions from *K* different source populations (clusters) that are inferred from the data. Since we aimed at detecting genetic admixture between wild individuals and two different domestic strains, ADMIXTURE was run separately for each the three wild populations (Mare, Gravezon and Orb), along with the Atlantic and Mediterranean domestic strains, with a *K* value of 3. Bar plots of estimated ancestry proportions given by the *Q*-values estimates by ADMIXTURE were produced with R v. 3.4.3. (Team, 2015).

We used the NEWHYBRIDS v2.0 software (Anderson and Thompson, 2002) to assign individual genotypes to Atlantic and Mediterranean parental lineages and different hybrid pedigrees (including F1 and F2 hybrids, and first generation backcrosses (BC) in each direction) in each of the three rivers. Domestic Atlantic individuals (N = 54) and wild individuals with a Mediterranean ancestry greater than 95% (N = 29 individuals: N_Mare_ = 4, N_Orb_ = 10, N_Gravezon_ = 15; Table 1) were considered as pure parental references for the Atlantic and Mediterranean lineages, respectively, using parental priors with the *z* and *s* options. We only used the most informative SNPs between these parental reference populations to increase the detection efficiency of hybrid categories. In total, 196 SNPs with a Weir and Cockerham’s *F*_ST_ > 0.85 were retained to run NEWHYBRIDS using 50,000 iterations after 100,000 burn-in steps.

### Inference of local ancestry

The inference of local ancestry was performed with ELAI v1.01 (Guan, 2014), which relies on a two-layer hidden Markov model to detect the structure of haplotypes in unrelated individuals. The program was run separately for each of the 40 linkage groups (LG) of the *S. trutta* linkage map (Leitwein *et al.*, 2017). We took advantage of the strong collinearity between the *Salmo trutta* and *S. salar* genomes (Leitwein *et al.*, 2017) to anchor the 4,000 mapped RAD loci of the brown trout linkage map onto the reference genome of *S. salar.* This strategy allowed us to determine the relative mapping positions of a large number of additional RAD loci that were not present on the brown trout linkage map, using their relative positions on the Atlantic salmon reference genome. A list of ordered RAD loci was created for each of the 40 brown trout LGs, and passed to the STACKS populations module using the whitelist option (-W) with the previously described filters to generate an ELAI input file for each brown trout LG. ELAI was run separately for each wild Mediterranean population. In each run, we used the 54 Atlantic and the 40 Mediterranean domestic individuals as source populations, and also used wild individuals with a Mediterranean ancestry greater than 95% (N = 29; Table 1) as a third source population. The number of upper clusters (-C) was set to 3 (i.e. because wild fish potentially originated from up to three source populations: domestic Atlantic, domestic Mediterranean and wild Mediterranean), the number of lower clusters (-c) to 15 (i.e. 5C, as in Guan 2014), the number of admixture generations (-mg) to 10 (i.e. approximately corresponding to the beginning of stocking), and the number of expectation-maximization steps (-s) to 20. The ancestral allele dosage from a given source population (see below) at each SNP was finally plotted for each individual along each LG to generate ancestry profiles with R (Team, 2014).

### Estimation of tract length and chromosomal ancestry imbalance

The junctions between haplotypes originating from different source populations were identified from the analysis of individual ancestry profiles produced by ELAI, as illustrated in Figure 1. At each variable position, we conservatively considered as evidence for 0, 1 or 2 haplotype copies from a given source when the inferred ancestry dosage was within the range [0, 0.05], [0.95, 1.05], or [1.95, 2], respectively (Figure 1A). Therefore, the junctions between haplotypes from different sources occurred within ‘uncertainty areas’ where the ancestry dosage lays between 0.05 and 0.95, or between 1.05 and 1.95. For each ‘uncertainty area’ within each LG of each individual, the junction between the ending and starting positions of two haplotypes from different sources was determined as the position where the estimated ancestry-dosage curve produced by ELAI crossed the 0.5 or 1.5 values (Figure 1A).

**Figure 1.**
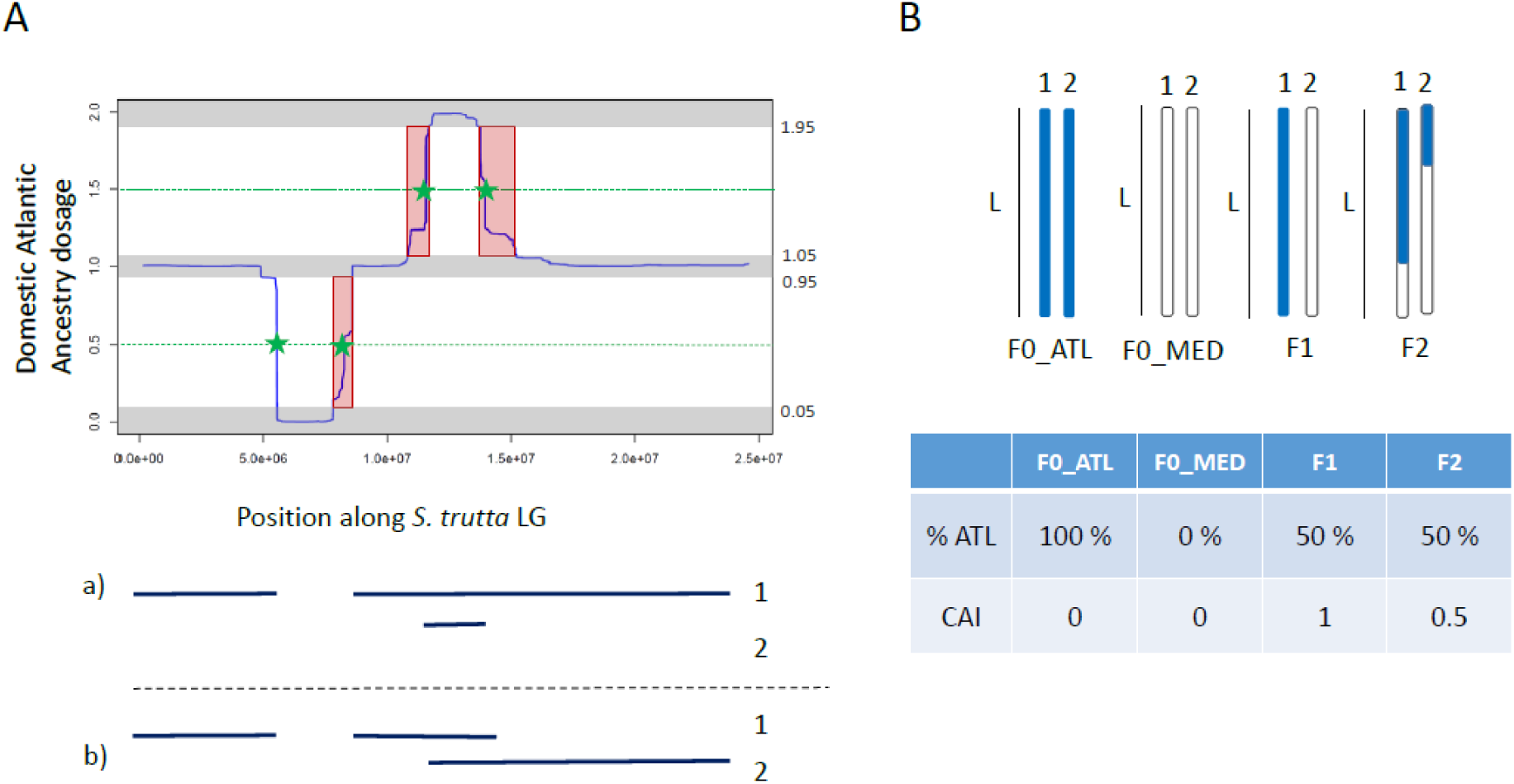
Detection of the haplotypic origin along each LG of brown trout with ELAI (Guan, 2014). The left panel (A) represent one individual output provided by ELAI, in which the ancestry dosage (0, 1, 2) of the Atlantic haplotype is represented as a function of the position along Salmo trutta LG in bp. Horizontal grey zone represent the 0.05 confidence interval of ancestry dosage. Green stars represent the junctions between the ending and the starting position of each Atlantic haplotype. Red boxes represent the uncertainty areas (see text for details). On this left panel, (a) and (b) represent the two concurrent interpretations of haplotypic structure along each homologue (1 and 2). Each interpretation satisfies the description of variation in Atlantic allelic dosage presented above. The right panel (B) describes the theoretical expectations of the percentage of domestic Atlantic haplotypes and the chromosome ancestry imbalance (CAI) for: F0_ATL: pure domestic Atlantic homologues with the Atlantic ancestry represented in blue, F0_MED: wild Mediterranean, F1: hybrids resulting from a crossing between two pure parental population (F0_MED and F0_ATL), and F2: theoretical hybrids resulting from a crossing between two F1 individuals. The percentage of Atlantic ancestry of each individual represent the sum of all Atlantic haplotypes length divided by the chromosomal length (L). The CAI represent the difference of cumulated Atlantic haplotype length between the two parental homologues (1 and 2) divided by the haploid chromosomal length (L).

Because the approach implemented here with ELAI does not use phased haplotype data, the delimitation of tract junctions could not be unambiguously resolved in the particular case when the ancestry dosage was found to vary from 1 to 2 and back to 1 haplotype copy (Figure 1A). Such profiles may correspond to two alternatives situations, one involving the co-occurrence of a short haplotype inherited from one parent, fully overlapping a long haplotype inherited from the other parent (case a) in Figure 1A), and the second involving the presence of two medium-sized and partially overlapping haplotypes (case b) in Figure 1A). Although it is not possible distinguishing between these two alternatives, we systematically resolved these cases by considering the existence of one short and one long haplotype, in order to avoid over-splitting real long haplotypes in early-generation hybrids, which would artificially downwardly bias our estimates of chromosomal ancestry imbalance (see below). This choice is also consistent with the know history of uninterrupted stocking practices over the last decades, generating combinations of short (i.e. more ancient) and long (i.e. more recent) haplotypes originating from the same source within individual genotypes.

The identification of tract junctions allowed retrieving the number and length of tracts originating from each of the three source populations for each of the 40 LGs of each individual. Haplotypes lengths were then summed per origin across LGs and divided by the diploid genome size to estimate individual ancestry proportions from domestic Atlantic, domestic Mediterranean and wild Mediterranean sources. In order to evaluate the consistency of admixture proportions estimated using SNP frequencies and haplotype lengths, we tested the correlation between the levels of admixture estimated with ADMIXTURE and ELAI using Spearman test in R (Team, 2014). We further tested whether the percentage of individual domestic ancestry from each domestic source correlates with the genome-wide averaged heterozygosity computed with VCFTOOLS v0.1.14 (Danecek *et al.*, 2011).

### Quantifying the timing of hybridization and introgression

In order to characterize the dilution of Atlantic alleles within Mediterranean genomes over time, we defined the chromosomal ancestry imbalance (CAI, Figure 1B). For a given chromosome, the CAI represents the difference between the cumulated lengths of Atlantic haplotypes from each of the two parental homologues, divided by the haploid chromosome length. Therefore, the CAI ranges from 0 to 1, and can be averaged across the 40 LGs to provide a genome-wide measure for each individual. The CAI theoretically reaches its maximal value of 1 for F1 hybrids obtained by crossing two pure (i.e. non-admixed) parental populations, and is progressively reduced at every generation post-admixture due to recombination and random transmission of homologues from one generation to the next. Therefore, the CAI converges to 0 after a sufficient number of generations post admixture. The value of 0 also characterizes pure individuals (e.g., F0_ATL and F0_MED, Figure 1B).

The length distribution of tracts originating from the two domestic sources was used to estimate the timing of hybridization in each of the 3 wild populations. Following (Racimo *et al.*, 2015), the relationship between the time since admixture (*T* in generations) and the mean length of introgressed tracts (*l*) is given by:

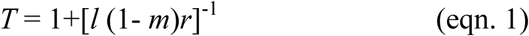

where *m* is the proportion of domestic ancestry within the recipient population, and *r* the recombination rate (in Morgan per base pair per generation). We used the genome-wide average recombination rate estimated in Leitwein *et al.* (2017), taking into account its variation across the genome to account for uncertainty. The mean tract length (*l*) in eqn. 1 was replaced by the *mode* of the length distribution of domestic tracts (either of Atlantic or Mediterranean origin). The rationale behind that choice was that the mode of the distribution better reflects the average time when the intensity of stocking was the most important, whereas the mean of the distribution would instead capture the presence of long tracts in early generation hybrids (Figure S1). Thus, using the mode of the distribution for estimating the timing of hybridization was considered more realistic to obtain conservation relevant estimates in this study. Individuals identified as ‘pure’ F0 individuals were discarded prior to building the length distributions of Atlantic and Mediterranean domestic tracts.

## Results

### SNP calling

A total of 1.5 billion of demultiplexed raw reads resulting in an average of 8.76 million reads per individuals were retained for the STACKS analysis. The previously established RAD catalogue (Leitwein *et al.* 2016) based on 64 individuals was updated using the 176 individuals considered in this study. After applying quality and population filters, we finally retained for subsequent analyses a total of 86,175 SNPs from 45,435 RAD loci, with a mean coverage depth of 47.7±17.9X per SNP per individuals and a percentage of missing genotypes of 4.20±2.9% per individuals.

### Clustering analysis

The ADMIXTURE program identified the three expected groups (*K* = 3) for each run performed separately for each wild population, including the domestic Atlantic and Mediterranean strains along with the wild populations considered in each run (either Gravezon, Orb or Mare; Figure 2). In the Gravezon River, 6 individuals were assigned as pure domestic Atlantic fish, and several individuals found with a 50% domestic Atlantic and a 50% wild Mediterranean ancestry were probably F1 hybrids. In the Orb River, the program found overall low proportions of domestic ancestries from either Atlantic or Mediterranean strains. In the Mare River, four individuals had more than 50% of Mediterranean domestic ancestry, and no putative Atlantic/Mediterranean F1 individual was detected, as in the Orb River. Moreover, each of the three ADMIXTURE runs revealed that the Atlantic domestic strain displays a low proportion of Mediterranean ancestry, with consistent estimates of individual admixture proportions among runs (Figure 2).

**Figure 2.**
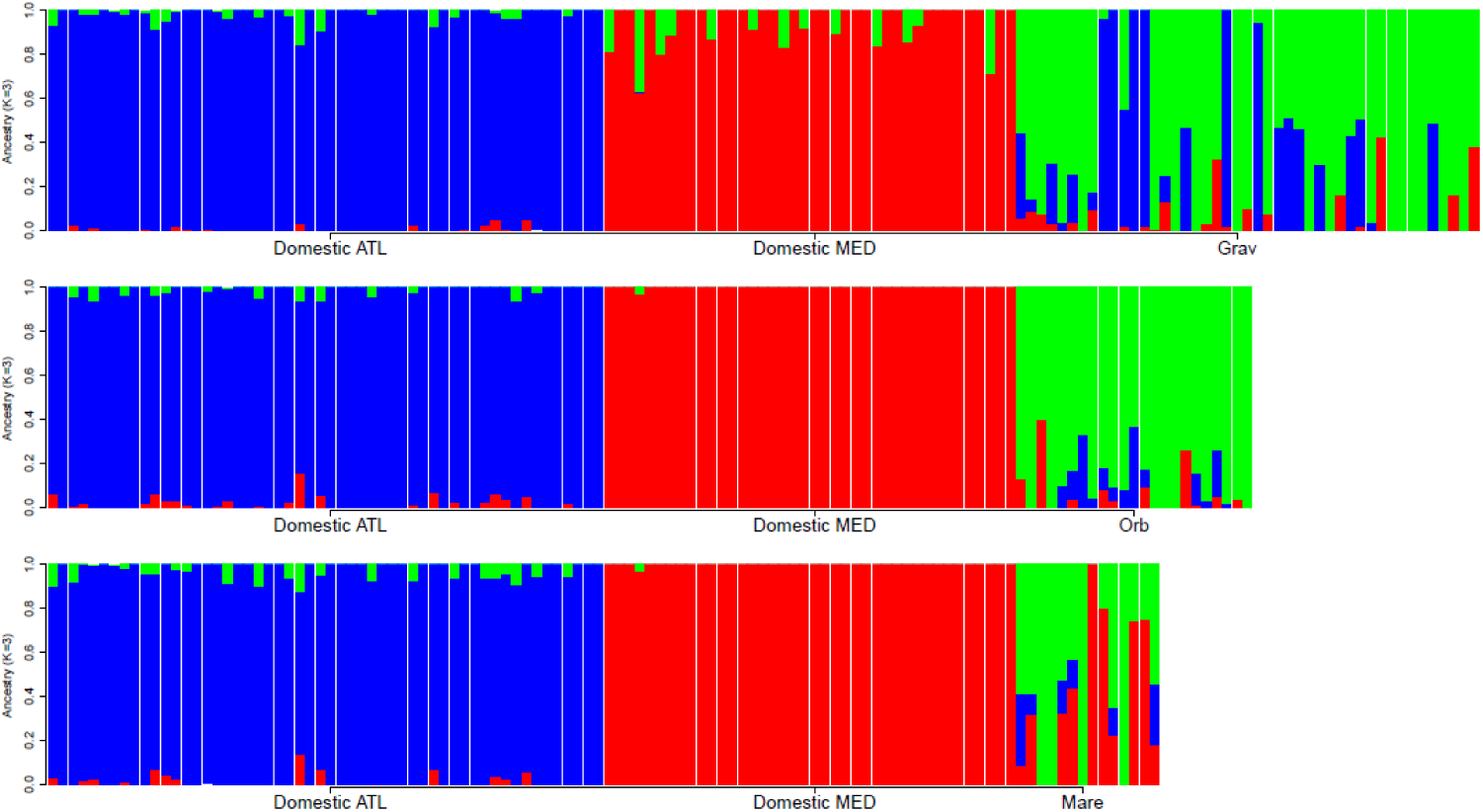
Plots of the individual ancestry inference for the 86,175 SNPs present in wild-caught brown trout. Analyses were separately run (K = 3) for individuals of the three wild populations (from top to bottom: Gravezon; Orb and Mare populations) using ADMIXTURE v1.3. Each individual is represented by a single vertical line, with colors indicating the estimated ancestry in each of the K = 3 groups: Atlantic domestic strain (blue), Mediterranean domestic strain (red), or wild Mediterranean (green).

The numbers of pure, F1, F2 and first generation backcrosses between Atlantic and Mediterranean lineages present in each wild population were estimated with NEWHYBRIDS. Despite setting priors on parental samples, two Atlantic individuals from the hatchery were not assigned to the parental Atlantic lineage, indicating possible admixture in their recent ancestry (Table S1). Similarly, two individuals from the Orb, one from Gravezon and four from Mare River were not assigned as ‘pure’ wild individuals despite being defined as parental individuals in the prior settings (Table S1). According to NEWHYBRIDS, hybrid genotypes were present in all of the three wild populations, including seven F1’s in the Gravezon River, and four, three and five F2’s in the Orb, Gravezon and Mare rivers, respectively (all with ≥95% probability of assignment, Table S1). First generation backcross genotypes were mostly detected in the Orb River with NEWHYBRIDS (Table S1).

### Inference of local ancestry

Local ancestry inference was performed in ELAI using 75,684 mapped SNPs distributed along the forty *S. trutta* LGs for 53 admixed individuals captured in the wild (13 from the Orb, 30 from the Gravezon and 10 from the Mare; Table S2), using individuals identified as pure domestic (ATL-Dom or MED-Dom) or wild Mediterranean (MED-Wild) as reference samples. For clarity, Figure 3 provides four examples illustrating the inference of local ancestry profiles along a LG in individuals with different ancestries. It first shows (Figure 3, upper left panel) an introgressed Atlantic haplotype of about 3 Mb (ATL-Dom ancestry dosage = 1) within a wild Mediterranean fish (MED-Wild ancestry dosage = 2). Then, the upper right panel describes a typical profile of a F1 hybrid resulting from the first generation of crossing between a pure domestic Atlantic and a pure wild Mediterranean individual, both contributing to an ancestry dosage of 1 all along the LG. The bottom left panel illustrates a F1 hybrid resulting from the crossing of a pure domestic Atlantic and a wild Mediterranean parent introgressed by the Atlantic domestic strain (i.e. resulting in homozygous Atlantic positions with an ATL-Dom ancestry dosage = 2). The last panel illustrates the typical profile of a pure domestic Atlantic individual (ATL-Dom ancestry dosage = 2). In these four selected examples, no signature of domestic Mediterranean introgression was observed (MED-Dom ancestry dosage = 0).

**Figure 3.**
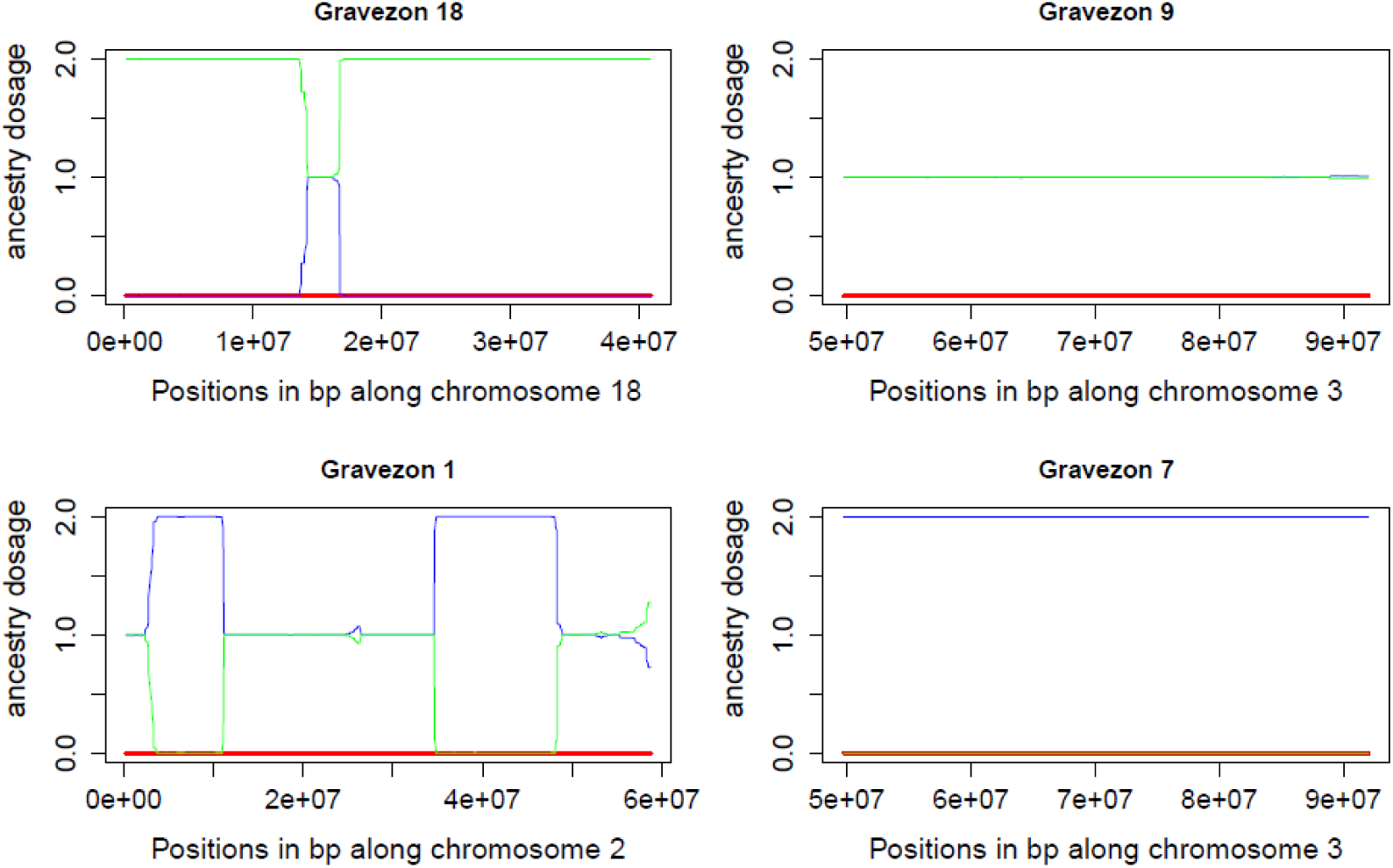
Examples of four individual plots of the local ancestry inference run established with ELAI (Guan, 2014). The ancestry dosage (y-axis; see Fig. 1A) is represented along LGs 4, 3 and 2 (x-axis), for four distinct Gravezon individuals (18, 9, 1 and 7). In green: the wild Mediterranean ancestry, in red: the domestic Mediterranean ancestry and in blue: the Atlantic domestic ancestry.

Junctions between adjacent ancestry tracts were generally represented by sharp transitions in chromosomal ancestry profiles. On average, regions of uncertain ancestry represented a 6.54±5.75% fraction of the genome around domestic Atlantic tracts and a 12.15±8.15% fraction around domestic Mediterranean tracts. This result indicates a reasonably high precision to identify the junctions between adjacent ancestry tracts.

### Analysis of hybridization and admixture from local ancestry tracts

Local ancestry patterns were summarized across LGs to determine the total number and the cumulative length of introgressed haplotypes along the genome of each individual, in order to provide an estimate of the genome-wide ancestry from each source population. The number and the mean length of introgressed haplotypes of domestic Atlantic and Mediterranean ancestry were found to considerably vary among the 53 admixed individuals considered in this study (Table S2). Within individuals, the length of ancestry tracts was also highly variable (Table S2), partly because chromosome length (which varies among chromosomes) influences the length of tracts during the first generations of admixture. Individual admixture proportions calculated as the percentages of Atlantic and Mediterranean domestic tracts relative to the total length of chromosomes were found to be significantly positively correlated to the percentages of domestic ancestry computed with ADMIXTURE (ATL-Dom: *rho*_Spearman_= 0.95, *p* < 2.2e-16; MED-Dom: *rho*_Spearman_= 0.88, *p* < 2.2e-16, Figure 4 A and B). This result confirms that the genome-wide average ancestry estimated with ELAI is highly consistent with the estimates obtained using the more classical ADMIXTURE approach, despite the existence of a small fraction of ‘uncertainty areas’ where local ancestry could not be determined. However, ADMIXTURE reported the existence of individuals with no domestic ancestry – either from Atlantic or Mediterranean strains – that were found with small but non-negligible domestic ancestries in ELAI (Figure 4). This discrepancy in estimating the presence/absence of non-admixed individuals in wild populations may be explained by the tendency of ADMIXTURE to minimize the estimated admixture fraction of the least introgressed samples.

**Figure 4.**
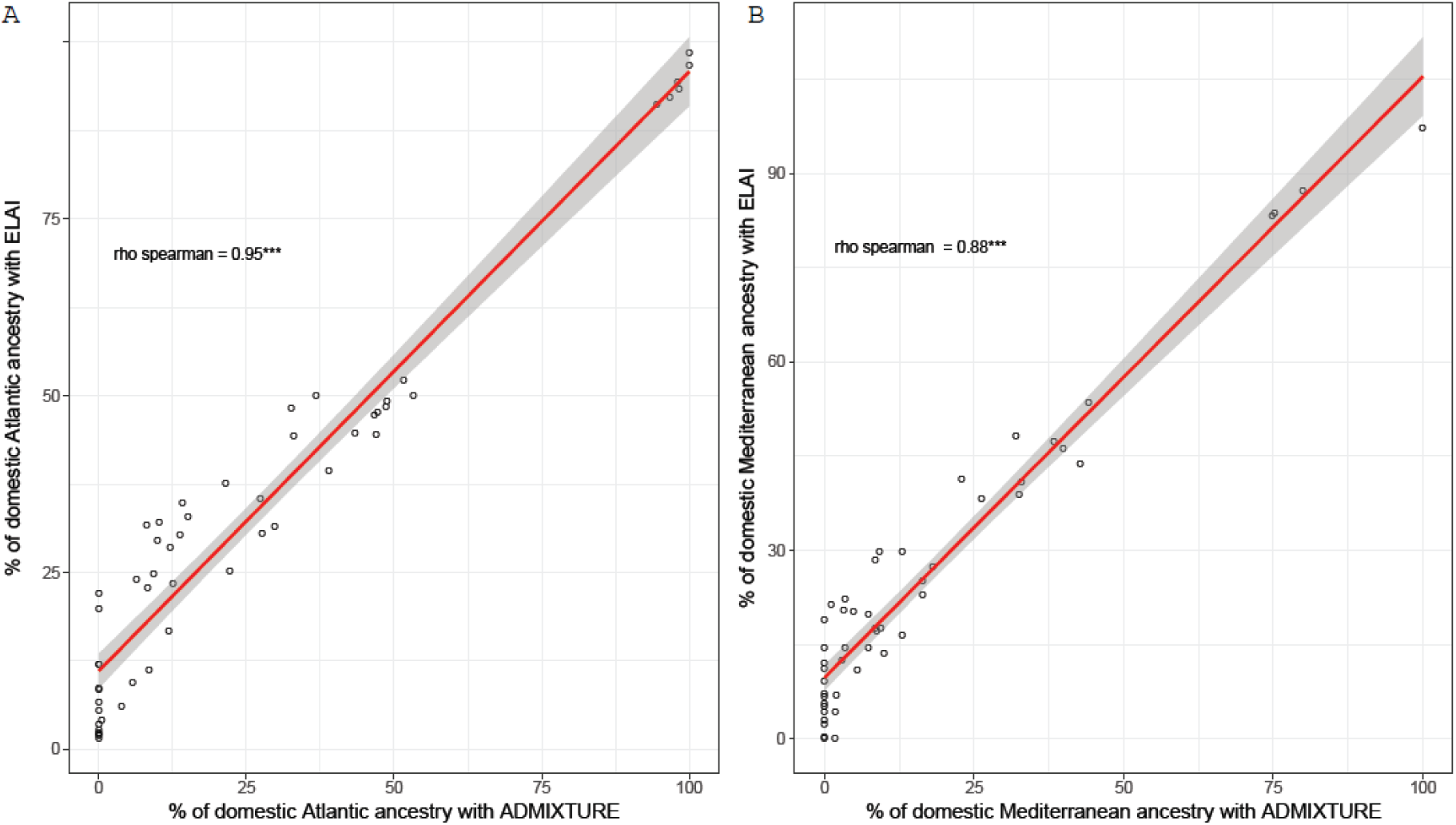
Positive correlation between the percentage of domestic Atlantic (A) and Mediterranean (B) ancestry computed with ELAI and ADMIXTURE (rho _spearman_= 0.95 and 0.88, p-value <2.2e-16, respectively)

We then compared local ancestry patterns among wild individuals in order to characterize the diversity of admixed genotypes between wild Mediterranean populations and the more evolutionary distant Atlantic strain. The estimated chromosomal ancestry imbalance (CAI) was expressed as a function of the percentage of Atlantic ancestry estimated with ELAI in a triangle plot, together with the number of Atlantic haplotypes (Figure 5). These estimates were compared to the theoretical expectations that the maximum CAI=1 for F1 hybrids, and CAI=0 for pure Atlantic (ATL-Dom) and Mediterranean (MED-Wild) individuals. Moreover, the expected number of Atlantic haplotype tracts in pure Atlantic individuals corresponds to the diploid chromosome number (2n=80), whereas it is equal to 0 in pure Mediterranean fish. These theoretical expectations were not totally met by real F1 hybrids and parental samples (Figure 5), indicating that introgression has occurred in both parental populations (ATL-Dom and Med-Wild). This was further supported by the results obtained with NEWHYBRIDS based on 196 SNPs with *F*_ST_ > 0.85. Indeed, the parental Atlantic samples identified by NEWHYBRIDS clustered close to the theoretical position of pure Atlantic parents in the triangle plot, although they contained more than 80 Atlantic tracts and displayed small fractions of Mediterranean ancestry (Figure 5). Likewise, fish identified as F1 showed a reduced CAI and an increased number of Atlantic ancestry tracts compared to what was expected for F1 hybrids produced between non-introgressed parental populations. Discrepancies were also found to occur for some individuals beyond first hybrid generation. For example, F2 individuals should have similar Atlantic ancestry to F1 hybrids, but an estimated CAI of ∼0.5 vs ∼1.0. Four individuals were found to meet these expectations in Figure 5 (i.e. individuals with 72, 87, 97 and 98 Atlantic haplotypes located in the center of the triangle plot), three of which have been assigned as F2 with NEWHYBRIDS. The NEWHYBRIDS analysis further identified several F2 in the Mare and the Orb rivers (Figure 5), which did not present either a number of Atlantic haplotypes or an Atlantic ancestry compatible with the expectation of the F2 category (Figure 5, Table S2). The finding of reduced Atlantic tract numbers and Atlantic ancestries (down to 55 and 25%, respectively) suggested that these individuals were more likely backcross genotypes produced between a F2 and a backcross parent. Moreover, two individuals were assigned as pure wild Mediterranean fish with NEWHYBRIDS (Table S2), while they appeared to be more likely backcrosses produced between a F1 and a Mediterranean parent. Indeed, one of them displayed a CAI of 0.49 together with Atlantic ancestry of 25.3%, which corresponds to what is expected for a first generation backcross. Therefore, our results probably illustrate some limitation of NEWHYBRIDS to assign genotypes to predefined hybrid classes in the presence of complex admixtures that go beyond the categories that are specified prior to the analysis. However, both methods produced mostly similar results for parental, F1 and F2 classes.

**Figure 5.**
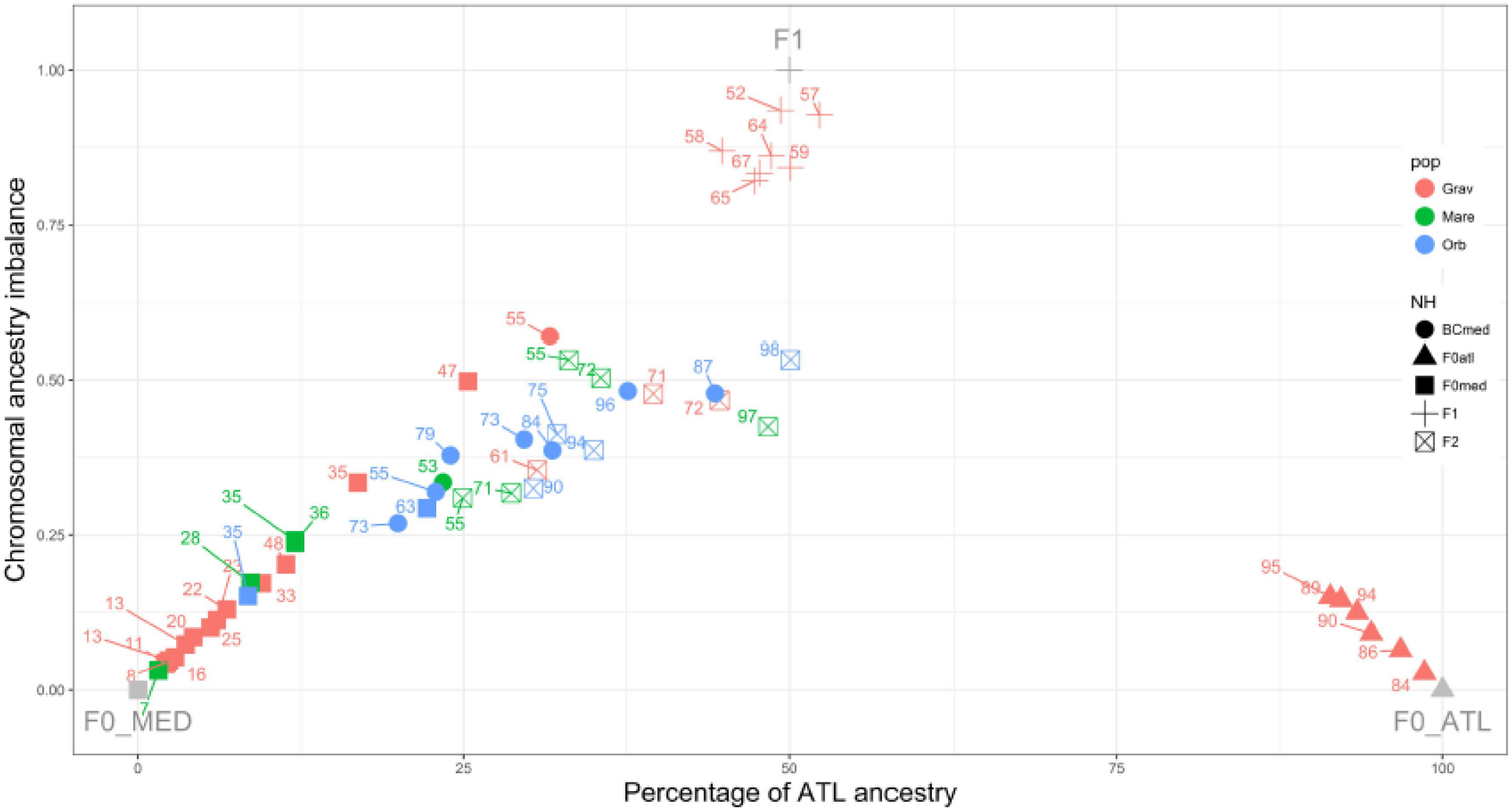
Plot of the chromosomal ancestry imbalance in function of the percentage of Atlantic ancestry for each wild-caught admixed individual considered. Grey points represent the theoretical expectations for F0_MED (without Atlantic ancestry, thus a CAI of 0 because individuals are theoretically pure Mediterranean), F0_ATL (without wild Mediterranean ancestry, thus a CAI of 0 because individuals are theoretically pure Atlantic) and F1 individuals (with half Atlantic ancestry and half wild Mediterranean ancestry, thus a CAI of 1 because the chromosomal ancestry imbalance is maximum between homologues). The colors represent the three wild populations: red for wild-caught individuals of the Gravezon River, then green for the Mare and blue for the Orb individuals, respectively. The different points shape represent the hybrids categories as obtained with NewHybrids. The estimated numbers of Atlantic haplotypic tracts for each individual are reported and ranged from 7 to 98 with theoretical expectation ranging from 0 to 80.

The analysis of CAI, estimated percentage of Atlantic ancestry, and estimated number of Atlantic tracts in wild individuals showed a complete absence of backcross individuals in the Atlantic genetic background (Figure 5). On the contrary, we found extensive backcrossing in the opposite direction, with individuals showing CAI and Mediterranean ancestry values compatible with a range of backcross pedigree beyond the first backcross generation (Figure 5). This was illustrated by a decreasing gradient in CAI, Atlantic ancestry and estimated number of Atlantic tracts between F2 hybrids and the least introgressed Mediterranean individuals.

The percentage of domestic ancestry was also compared to the individual heterozygosity in order to evaluate the consequences of admixture and introgression on the level of polymorphism. We found a significant positive correlation between the percentage of domestic Atlantic ancestry and individual heterozygosity (Figure 6A, *rho*_Spearman_ = 0.88, *p* <2.2e-16). Conversely, a significantly negative correlation was found between the percentage of domestic Mediterranean ancestry and individual heterosigosity (Figure 6B, *r*_Spearman_ = −0.44, *p* <2.2e-16).

**Figure 6.**
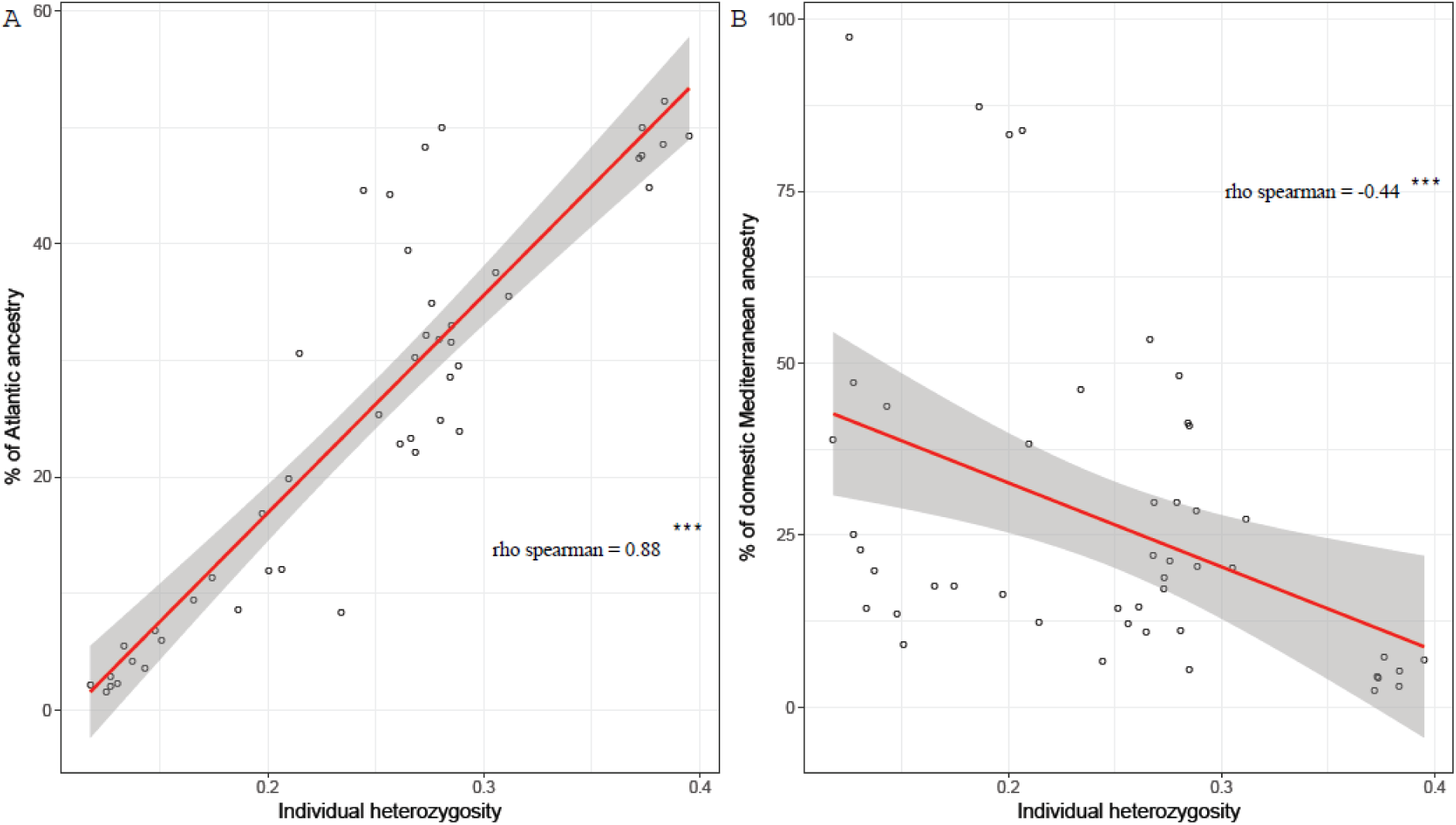
Correlations between the percentage of domestic ancestry and the individual heterozygosity. (A) Positive correlation between the percentage of domestic Atlantic ancestry (rho_spearman_=0.88, p-value<2.2e-16). (B) Negative correlation between the percentage of domestic Mediterranean ancestry and the individual heterozygosity (rho_spearman_=-0.44, p-value<2.2e-16)

### Estimation of time since admixture

In order to estimate the time in generations since the introduction of Atlantic and Mediterranean domestic haplotypes within each wild Mediterranean population, the mode of : the length distribution of domestic haplotypes was used within each population. The individuals formerly recognized as “pure” domestic Atlantic (Figure 5, Gravezon’s
individuals at the right corner) were removed from data before estimating *T*, as well as four individuals with more than 80% of domestic Mediterranean ancestry identified in the Mare River (Table S2). The proportions of Atlantic ancestry in each wild population were relatively close from each other (0.244, 0.228 and 0.299 for the Gravezon, the Mare and the Orb rivers, respectively). The modes of the estimated length of Atlantic haplotypes were more heterogeneous across populations (6,533,308 bp, 7,878,364 bp and 5,421,128 bp for the Gravezon, the Mare and the Orb rivers, respectively) (Figure S2). Considering that the mean recombination rate was estimated to 0.88cM/Mb in brown trout (Leitwein *et al.* 2017), and taking in account the recombination rate variation by using the first and the third quartile estimates of the recombination rate (0.37 cM/Mb and 1.13 cM/Mb, respectively; derived from Leitwein *et al.*, 2017). The time *T* was estimated to 24.01 generations ([18.92-55.73] generations using the 1^st^ and 3^rd^ quartiles of the estimated recombination rate) for Gravezon, 19.70 generations ([15.55-45.46] generations) for the Mare and 30.90 generations ([24.28-72.10] generations) for the Orb (Figure 7)

**Figure 7.**
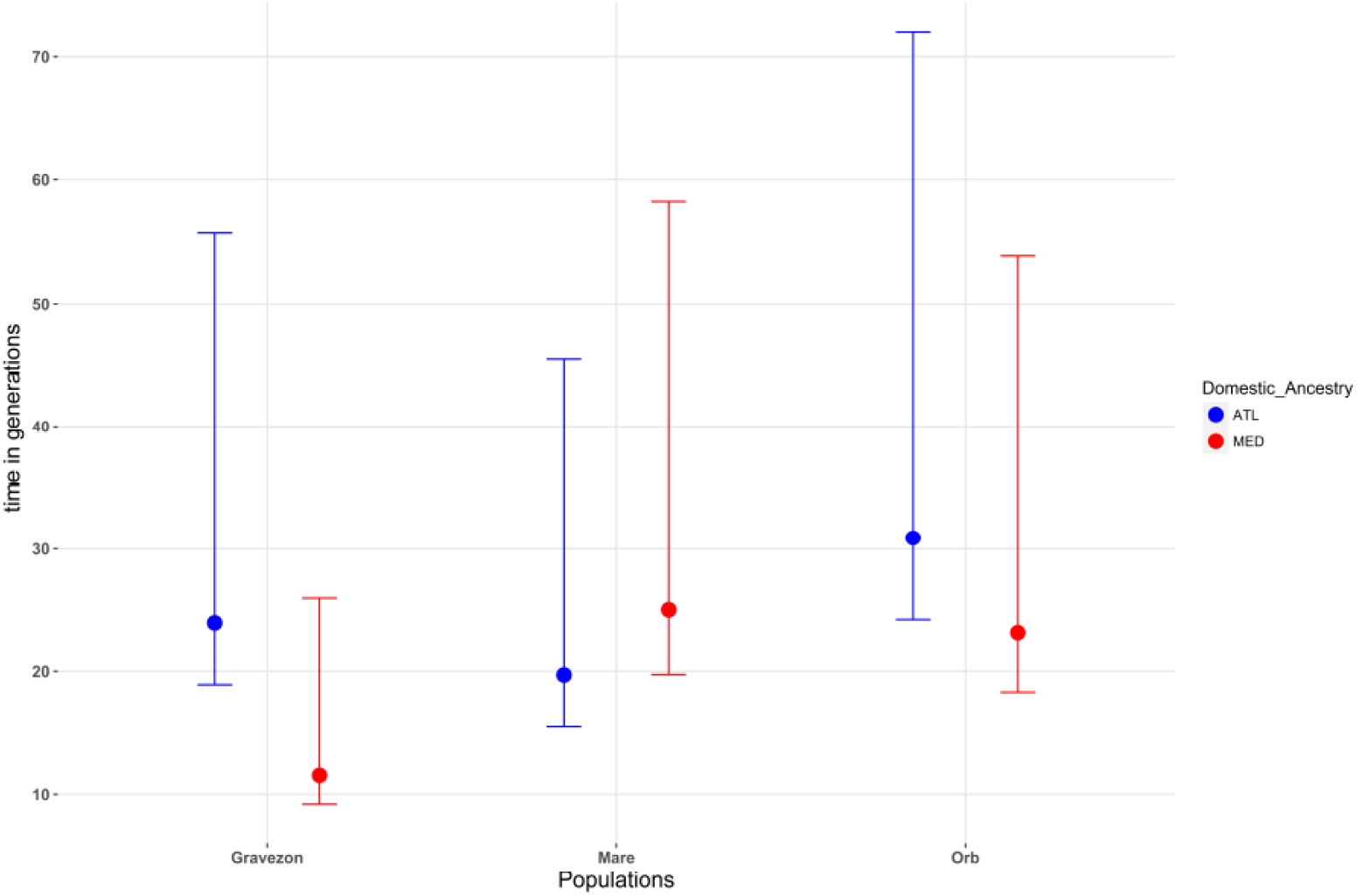
Estimates of the number of generations (t) since introgression of both Atlantic (in blue) and Mediterranean (in red) domestic strains in each wild Mediterranean populations. The confidence intervals have been estimated using the first and third quartiles of recombination rate estimates provided by Leitwein et al. (2017). See text for details.

Similarly, we estimated *T* for domestic Mediterranean admixture into the wild populations. The proportions of Mediterranean ancestry in the populations were 0.155, 0.381 and 0.242 for the Gravezon, the Mare and the Orb rivers, respectively. The mean lengths of domestic Mediterranean haplotypes were 12,786,692 bp, 7,633,631 bp and 6,740,239 bp for the Gravezon, the Mare and the Orb rivers, respectively (Figure S2). *T* was estimated to 11.51 generations ([9.19-26.01] generations) for Gravezon, 25.07 generations ([19.75-58.25] generations) for the Mare and 23.23 generations ([18.31-53.87] generations) for the Orb (Figure 7).

## Discussion

By combining Anderson and Stebbins’ (1954) ‘genic blocks’ view with population genomic approaches to characterize a complex three-way admixture, we here provide an improved description of the genome-wide consequences of human-mediated hybridization in wild populations of Mediterranean brown trout. In Southern France, as in many other locations throughout Europe (see Bohling *et al.* 2016), domestic strains derived from different brown trout evolutionary lineages (Atlantic and Mediterranean; Bernatchez, 2001; Sanz, 2018) were commonly used for supplementation. The Atlantic domestic trout has been used for decades to enhance wild Mediterranean populations, but since 2004, this strain was largely replaced by a local domestic Mediterranean strain developed using wild breeders from the Gravezon River, for local supplementation. In this study, RAD-derived SNPs were used to provide a genome-wide picture of individual admixture within each of three wild Mediterranean populations. We performed local ancestry inference in each individual using 75,684 mapped SNPs without using phase information, and evaluated the performance of this approach to estimate genome-wide average ancestry compared to classical ancestry inference methods. We also compared the percentage of Atlantic ancestry and the chromosomal imbalance for Atlantic ancestry between homologues across different categories of parental, admixed and introgressed genotypes. Finally, we used the length distribution of introgressed domestic haplotypes to estimate the time since the maximum rate of admixture with both the Atlantic and Mediterranean domestic strains in each local population. We show that admixture had dramatically different consequences on the level of polymorphism depending on the domestic source population used for stocking, thus providing important information to future conservation actions.

### Tree-way admixture inference

Individual proportions of domestic Atlantic, domestic Mediterranean and wild Mediterranean ancestry were assessed with ADMIXTURE and ELAI using 86,175 SNPs and 75,684 mapped SNPs, respectively (Figure 2; Table S2). Contrary to microsatellites data that did not allow the detection of the domestic Mediterranean strain in Gravezon river (Berrebi and Schikorski, 2017), admixture from the two domestic strains (i.e. Atlantic and Mediterranean) was efficiently detected in each river using the dense SNP marker dataset.

The individual proportions of domestic and wild ancestry assessed with ELAI using the haplotypic ancestry information showed a high consistency with the ancestry proportions inferred with ADMIXTURE. Indeed, a strong positive correlation was observed between ADMIXTURE and ELAI results for both Atlantic and Mediterranean domestic ancestry. Since the genome-wide ancestry proportions are consistent between methods for both domestic sources, it reinforces the value of the local ancestry inferences performed with ELAI, which has not yet been evaluated for its performance on similar data.

Both methods found six individuals sampled in the Gravezon River that were assigned as domestic Atlantic trout. Because the Hérault French Fishing Federation has stopped supplementing with the domestic Atlantic fish in this region, these individuals were probably escaped from a private hatchery during flooding events that are frequent in Mediterranean streams. Besides these six Atlantic fish, a low to moderate percentage (from 10 to 20%) of domestic Atlantic ancestry was detected in each of the three wild populations. This likely result from past introgression of Atlantic alleles which starting since the beginning of supplementation practices. Domestic Mediterranean ancestry was also detected in all three local populations, especially in the Mare River where four individuals over 14 had a high percentage of domestic Mediterranean ancestry (>60%), probably due to recent stockings in this river.

Our analyses also revealed discrepancies between methods, especially for estimating the lowest proportions of admixture within wild individuals. Indeed, ADMIXTURE assigned several individual as pure (i.e. zero percent of domestic ancestry, Figure 4) whereas ELAI detected low to moderate proportions of admixture (i.e. 0 to 20 %, Figure 4). This might reflect some limits in the inference of ancestry proportions with ADMIXTURE, which tends to minimize the level of admixture in the least introgressed genotypes. This discrepancy is also possibly explained by the fact that ADMIXTURE estimates ancestry proportions globally across the genome, whereas ELAI performs local ancestry inference. By taking advantage of the information contained in allelic association at linked markers, ELAI is expected to provide a finer scale picture of the mosaic of ancestry, which is supposed to improve the estimation of admixture proportions when introgression occurs over a small fraction of the genome. This interpretation, however, will need to be confirmed by a simulation study to evaluate the performance of ELAI in a similar context.

### Detection of Hybridization

The chromosomal ancestry imbalance (CAI) and the percentage of Atlantic ancestry estimated from ELAI output were used to characterize admixture between wild Mediterranean populations and the Atlantic domestic source. The percentage of domestic ancestry is equivalent to the hybrid index (Anderson, 1949; Buerkle, 2005), and the CAI to the interspecific heterozygosity, both classically used to infer hybridization patterns at an individual level (Larson *et al.*, 2013; Gompert and Buerkle, 2016; Wielstra *et al.*, 2017). However, by using haplotypes and the CAI, we were not limited by the use of only highly differentiated loci and thus were able to characterize admixture between domestic and wild Mediterranean populations exhibiting shallow differentiation. Moreover, the number and length distribution of introgressed tracts reflect the number of recombination events since hybridization, with a high number of short tracts revealing past introgression and a small number of long tracts revealing recent admixture (Gravel, 2012; Harris and Nielsen, 2013; Racimo *et al.*, 2015). These haplotype statistics are therefore more informative than the interspecific heterozygosity, which is not dependent of the number of generations since admixture beyond the first hybrid generation. For illustration, more abundant and longer Atlantic tracts were found in Mediterranean backcrosses compared to introgressed individuals displaying a low percentage of Atlantic ancestry, and showing less abundant and shorter Atlantic tracts (Figure 5). This result is consistent with a dilution of the Atlantic tracts over generations since the beginning of the supplementation history (i.e. supplementation with the domestic Atlantic strain has been stopped in this region). Moreover, the six ‘pure’ domestic Atlantic individuals which have been found in Gravezon River displayed a number of Atlantic haplotypes greater than the 80 expected from the diploid number of chromosomes (Leitwein *et al.*, 2017; Phillips and Ráb, 2001). Along with an estimated percentage of Atlantic ancestry lower than one hundred percent and a CAI greater than zero, this result indicates a possible admixture with Mediterranean individuals during the development of the Atlantic domestic strain. This hypothesis is further comforted by the low percentage of Mediterranean ancestry detected in the Atlantic domestic population with ADMIXTURE (Figure 2).

The use of haplotypic information provides a more detailed picture of recent hybridization and its consequences on admixture and introgression than NEWHYBRIDS (Anderson and Thompson, 2002), which can only assign individual genotypes to discrete predefined hybrid classes (e.g. typically parental, F1, F2, and first generation backcrosses). For example, several individuals in Gravezon, Mare and Orb rivers were assigned as pure Mediterranean by NEWHYBRIDS (F0med; Figure 5), while their percentage of Atlantic ancestry ranged from 8 to 32 %, with a number of Atlantic tracts ranging from 11 to 63 (Figure 5, Table S2). These individuals were unlikely to be pure Mediterranean, but rather introgressed individuals resulting from past hybridization with the domestic Atlantic strain at the beginning of supplementation. Several individuals were also assigned as F2 with NEWHYBRIDS while they were most likely a result of backcrossing, since their percentage of domestic Atlantic ancestry and their chromosomal ancestry imbalance were lower than expected for F2 individuals (Figure 5). This suggests that coupling haplotypic information to probabilistic model-based methods such as NEWHYBRIDS could greatly improve the detection of hybrids pedigree in a complex system like this one in the future.

### Timing of gene flow

The use of haplotype information can also allow modeling historical gene flow in order to date admixture events (Allendorf *et al.*, 2010; Gravel, 2012; Homburger *et al.*, 2015). Indeed, the length distribution of introgressed haplotypes within a population is a proxy of the time since introduction (Liang and Nielsen, 2014; Racimo *et al.*, 2015). Because introgressed haplotypes are progressively broken down into shorter fragments at each generation due to recombination events, long introgressed tracts denote recent admixtures while short haplotypes denote more ancient hybridization events (Racimo *et al.*, 2015). For estimating the number of generations since the most intensive introgression of domestic strain into each river, we choose to use the mode of the distribution of introgressed tracts length instead of the mean length. Thus this estimation was predominantly based on shorter fragments reflecting past supplementation. A bias toward the detection of short fragments might have occurred in our dataset due to the method used for the delimitation of tract junctions that could not be unambiguously resolved (see methods). Besides, the recombination rate is highly variable along the genome (Leitwein *et al.* 2017). Although we accounted for such variation using the first and third quartiles of the observed distribution of recombination rate, the estimated time in generation is not expected to be precise and remain an approximation that should be interpreted with caution. In Gravezon and Orb rivers, we found that the number of generations since the most extensive events of hybridization was greater for the domestic Atlantic strain than for the domestic Mediterranean strain, revealing a more ancient introgression of domestic Atlantic alleles which was expected from the known history of introduction. Conversely, in the Mare river, the introgression of domestic Atlantic alleles was found more recent than the introgression from the Mediterranean strain. This could denote a recent undocumented introgression of domestic Atlantic strain into the Mare river. The history of supplementation in the Orb watershed is not well documented, but according to stocking practices in the area, the use of the Atlantic domestic strain should be more ancient and should have stopped since 2004, as observed in the Gravezon and the Orb rivers. However several pure domestic Atlantic individuals have also been found in Gravezon river, and it is therefore possible that the Mare river has undergone an unknown recent event of domestic Atlantic admixture.

### Implications for brown trout conservation genomics

Genotype data at unlinked makers have been commonly used to assess hybridization and genome-wide levels of admixture and introgression for a large range of conservation genomics issues (Hohenlohe *et al.*, 2013; Hassanin, 2015; Lamaze *et al.* 2012; Bradbury *et al.* 2015; Le Moan *et al.* 2016; King *et al.*, 2015; Rougemont *et al.*, 2017). Nevertheless, the use of a linkage maps for assessing the relative order and linkage disequilibrium among loci, and the increasing availability of references genome allow to access haplotypic information which has the potential to increase the sensitivity of admixture estimates (Allendorf *et al.*, 2010), and allow to estimate the time since hybridization (Racimo *et al.*, 2015).

In this study, we took advantage of the linkage map available for the brown trout (Leitwein *et al.* 2017) and the Atlantic salmon reference genome (Lien *et al.* 2016), to assess introgression at a fine scale level in a complex system, where more than one domestic lineage has been used for supplementation. As a result, we were able to describe the complex pattern of admixture and introgression resulting from multiple genetic interactions between wild local Mediterranean populations and domestic Atlantic and Mediterranean strains (Leitwein *et al*., 2016). Our approach seems particularly relevant to identify allochtonous and autochtonous sources of supplementation for a conservation biology perspective, especially when local and domestic stains are genetically close to each other. In particular, when the development and prioritization of conservation actions is ruled by the presence of ‘pure’ wild individuals increasing the value of a stream or a watershed (Hansen and Mensberg 2009), it is often required to prioritize the identification of pure individuals in the wild. Nonetheless, we show in this study that the identification of pure individuals was sensitive to the method used and that no pure individuals likely remains in the studied populations, which is not surprising considering the history of supplementation. An inference relying on ADMIXTURE alone would have led to the wrong conclusion that domestic alleles have been totally purged through the time, which is not consistent with the finding of short domestic tracts in introgressed wild individuals. The complexity and the diversity of early-generation admixture have also been described here, without the need for using a priori on particular hybrids classes. Furthermore, the time since the most important event of supplementation from both domestic strains could have been estimated for each local population, which is meaningful for decision making in conservation.

Finally, our results showed dramatically different consequences of supplementation on the polymorphism of admixed populations, with opposite trends between the effect of introducing domestic Mediterranean or domestic Atlantic strains. Indeed, an increase in heterozygosity was associated to an increase in domestic Atlantic ancestry, as expected when mixing two distinct evolutionary lineages (Atlantic and Mediterranean), especially during first generations of hybridization (Allendorf *et al.*, 2012). This effect is here amplified by the fact that the Atlantic domestic strain is more genetically variable than the wild Mediterranean populations. On the contrary, an increase in domestic Mediterranean ancestry was associated to a decrease in heterozygosity (Figure 5), potentially indicating negative consequences associated with a loss of diversity and/or increased genetic load, as already suspected by Leitwein *et al.* (2016) because of the low polymorphism observed in the Mediterranean domestic strain. These observations suggest that the mosaic of individual haplotypic ancestry should be more thoroughly analyzed in order to better understand how hybridization/introgression impacts the genomic makeup and potentially the fitness of wild brown trout populations. An in-depth analysis of the genome-wide landscape of introgression is now needed to identify genomic regions where introgression is higher, or on the contrary, lower than expected under neutral introgression. Such approach will be highly informative for understanding the historical and the selective consequences of introgression (Sankararaman *et al.*, 2014; Racimo *et al.*, 2015). For example, we could detect signatures of adaptive introgression from the Atlantic domestic lineage into wild Mediterranean populations, or even, genomic regions that are resistant to gene flow because of negative selection against deleterious introgression. The detection of genomic region under positive or negative selection, will be of prime importance for future conservation and management actions (Allendorf *et al.*, 2001, 2010; Frankham, 2010; Stronen and Paquet, 2013; Garner *et al*., 2016).

## Acknowledgments

We thank E. Ravel and people from the Fédération Départementale de la Pêche de l’Hérault (France) and the Babeau hatchery for their help at several stages of this project. This project largely benefited from the Montpellier Bioinformatics Biodiversity cluster computing platform. Presequencing steps necessary for the production of SNP data were performed at the Genotyping and Sequencing facility of the LabEx CeMEB (Centre Méditerranéen pour l’Environnement et la Biodiversité, Montpellier), and RAD sequencing was performed at Montpellier GenomiX facility (Montpellier, France; http://www.mgx.cnrs.fr/). M.L. was partly supported by a grant from LabEx CeMEB.

## Data Accessibility

Supplemental Material, Table S1 contains NEWHYBRIDS results with sample ID and priors. Table S2 contains sample ID, the corresponding population, the mean percentage of Atlantic ancestry retrieved with ELAI and ADMIXTURE, the mean number and the mean length of introgressed Atlantic and domestic Mediterranenan haplotypes as well as the pedigree inferred with NEWHYBRIDS.

Raw demultiplexed sequence reads by individuals (fastq) are available at NCBI Short Read Archive under the study accession SRP136716 with one file per paired-end reads Sample.1.fq and Sample.2.fq and one file per unpaired reads Sample.rem.fq.

## Author Contributions

M.L and P.A.G conceived and designed the analyses. M.L performed the lab work, the analyses and wrote the manuscript. B.G and P.B supervised the research. All co-authors critically revised the manuscript and approved the final version to be published.

